# Lepidopteran prolegs are novel traits, not leg homologs

**DOI:** 10.1101/2022.06.30.498371

**Authors:** Yuji Matsuoka, Suriya Narayanan Murugesan, Anupama Prakash, Antónia Monteiro

## Abstract

Lepidopteran larvae have both thoracic legs and abdominal prolegs, yet it is unclear whether these are serial homologs. We examined the role of three Hox genes in proleg development in *Bicyclus anynana* butterflies using CRISPR-Cas9 and discovered that under a partial segment *abdominal-A* (*abd-A*) knockout, both types of appendages can develop in the same segment, arguing for prolegs being a novel trait, not a leg serial homolog. We also discover that *specificity protein (sp)* genes do not co-localize with Distal-less (Dll) in prolegs, as they do in legs, and that the proleg gene-regulatory network (GRN) mostly resembles the head-horn GRN, another novel trait in the lepidoptera. We propose that larval prolegs evolved from the co-option of a partial limb GRN into novel embryonic coordinates in the abdomen.

**One-Sentence Summary:** The chubby legs of caterpillars are evolutionary innovations

## Main Text

While most adult insects have three pairs of walking legs, juveniles display tremendous variation in both thoracic and abdominal appendages, only some of which are used for walking (*1, 2*). The origin, development, and evolution of such diversity in appendage number in insects, however, is still poorly understood. It is possible that in addition to processes of appendage repression and modification, from branchiopod-like ancestors, the origin of completely novel appendages might have taken place.

Previous research demonstrated that Hox genes regulate the number and type of abdominal larval appendages in some insect lineages. For instance, *Ultrabithorax (Ubx*) modifies the appendage primordia of the first abdominal segment (A1) of several insects into a transient embryonic glandular organ, the pleuropodium (*3, 4, 5*), that secretes a chitinase to facilitate embryonic hatching (*6*); and *abd-A* acts as a limb repressor in more posterior segments, in both *Tribolium* and *Drosophila* flies, partially via the repression of *Distal-less* (*Dll*) (*3, 7*). In Lepidoptera, however, a novel type of abdominal limb has evolved (Fig. 1A and B), and here *abd-A* is required for proleg development (*8, 9*). Furthermore, when *Ubx* is down-regulated together with overexpression of *abd-A* (due to a genomic mutation) prolegs develop in both A1 and A2 (*10*). These results indicate that *abd-A* is essential for proleg development and for repressing pleuropodium development, whereas *Ubx* is essential for pleuropodium development in A1 and for repressing proleg development in A2.

**Fig. 1.**
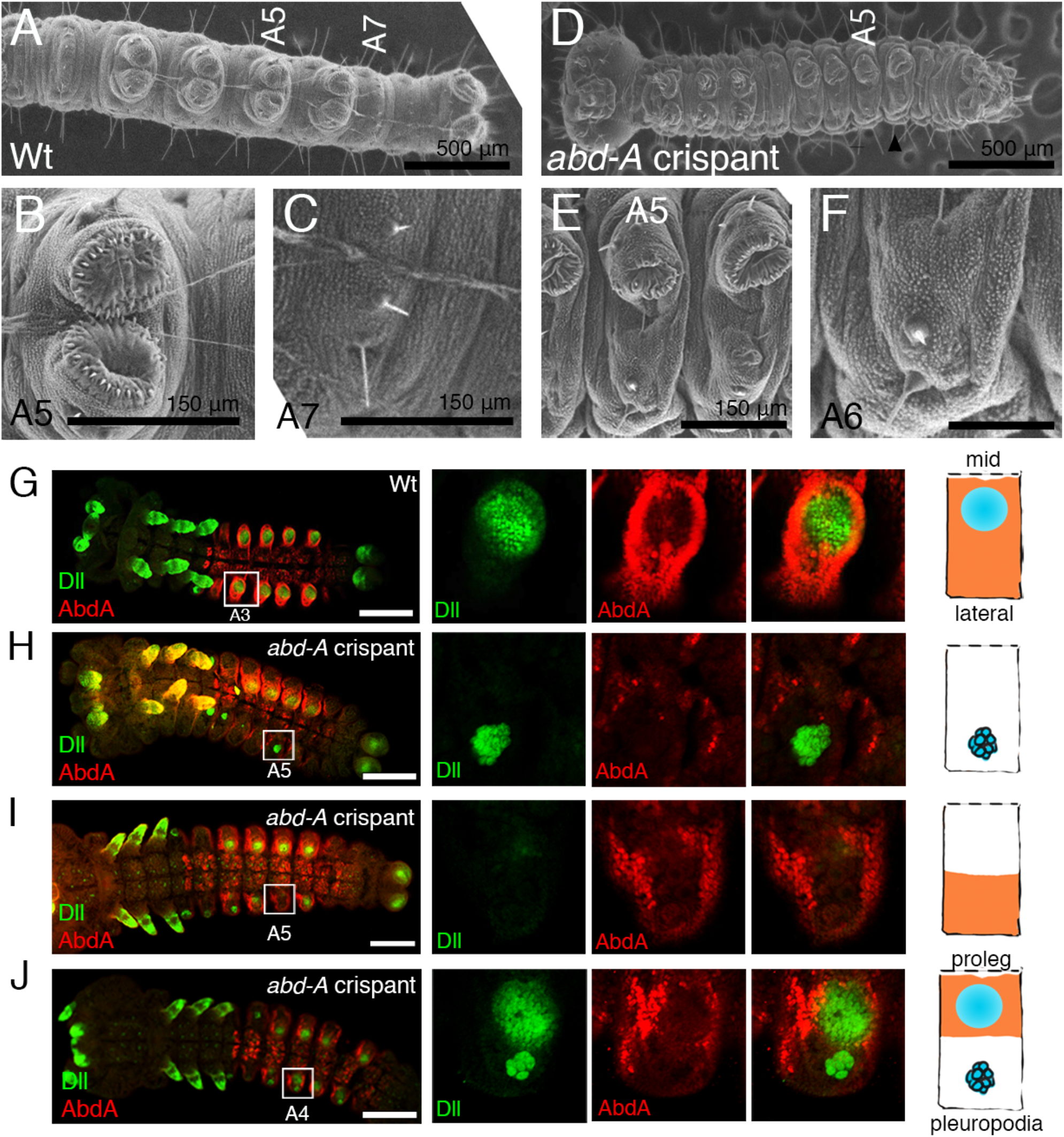
*abd-A* is necessary for proleg development and for repressing pleuropodia. (A-C) Wildtype prolegs (B) and limbless segment (C). (D-F) Prolegs are lost in *abd-A* mosaic crispants (E), and the segment shows features of the limbless segment (F). (G) Expression of Dll and Abd-A proteins in wildtype embryo. (H-J) *abd-A* crispants show three different types of Dll expression pattern due to mosaicism. Schematic diagram shows one side of the segment (midline of embryo is up). Expression domain of Abd-A is colored in orange.

These results show two instances of the same Hox gene having opposite effects on appendage development in different segments of one species. This is difficult to explain if these appendages are *bona fide* serial homologs. However, if thoracic legs and prolegs are different traits, using separate GRNs, each Hox gene might interact with each of the GRNs in a different way. *Ubx* could be both a modifier of the thoracic leg GRN, transforming it into pleuropodia in A1, and a repressor of the proleg GRN, preventing prolegs from developing in A2. Likewise, *abd-A* could be both a thoracic leg GRN repressor in A2 and more posterior segments, and an activator of the proleg GRN in those same segments. This can be tested under a partial segment Hox gene knock-out, where the two types of traits might be able to develop side-by-side in the same segment (*11*).

To test this dual GRN hypothesis, we first examined the detailed expression domains of three Hox proteins, Antennapedia (Antp), Ubx, Abd-A, as well as the limb development protein Dll, and then observed how CRISPR-Cas9 disruptions of each Hox gene affected thoracic and abdominal limb development, and the presence of Dll protein, during embryonic development.

Typical expression domains were found for all three proteins (Suppl. Fig. 1). Antp proteins were observed in the thoracic segments, Ubx proteins were observed in T3, and anterior compartments of A1 and A2 segments, and Abd-A proteins were present from the posterior part of A1 to A7, and part of A8 (Supp Fig. 1A, D, and G). These three Hox proteins were also present in the developing embryonic prolegs: Ubx in the more medial half-rim of the proleg tip, where Abd-A was absent (Supp Fig. 1C); Abd-A at the base of the proleg, anti-colocalized with Dll; Dll at the tip of the prolegs (Supp Fig. 1E and F); and Antp at the periphery of the proleg tips (Supp Fig. 1H and I). Most of these expression domains were previously observed in a different butterfly (*12*).

To test the function of the three Hox genes in limb development we used CRISPR-Cas9. Prolegs were not affected in *Antp* crispants, but thoracic legs were, in both conserved and novel ways relative to similar experiments in *B. mori* (see Supp Fig. 2, Suppl Text). *Ubx* crispant larvae displayed ectopic protuberances on the A1 and A2 segments that differed in morphology (see Supp Fig. 2). Those on A1 resembled thoracic legs because of the claw at the tip (Supp Fig. 2H and I), whereas those on A2 resembled prolegs with their row of crochets at the tip (Supp Fig. 2J and K). Prolegs were not visibly affected. *abd-A* crispants showed a similar phenotype as that observed in *Bombyx* RNAi embryos (*8, 9*). In mild cases, partial prolegs were retained, but in severe cases, *abd-A* crispants lost prolegs (Fig. 1E and F). These results indicate that *abd-A* is necessary for proleg development in *B. anynana*, as in *B. mori*.

To test the dual GRN hypothesis, we screened for *abd-A* mosaic mutants in embryos that only affected part of an abdominal segment. The effects of such disruptions must be examined in embryos as *abd-A* knockouts are expected to lead to ectopic pleuropodia (*3*), which will not protrude from the larval body wall, and can only be seen in embryos. To examine the possibility that prolegs are distinct traits from thoracic legs/pleuropodia we observed embryos using Dll immunostainings that mark these two traits in different ways - pleuropodia cells are fewer and larger than those of prolegs (Fig. 1H-J).

We observed a diversity of Dll protein patterns in *abd-A* mosaic crispants. In embryos where all the cells were lacking Abd-A in the same segment, Dll protein was lost from a proleg-epithelium pattern and acquired a pleuropodium-glandular pattern throughout the abdomen (Fig. 1H, Supp Fig. 3). In segments, where only the more medial region was lacking Abd-A, Dll protein expression in a proleg-pattern became patchy or disappeared altogether but pleuropodia were not visible (Fig. 1I, Supp Fig. 4). In segments where only the more lateral region was lacking Abd-A proteins, Dll expression was observed both in clusters of pleuropodium-style glandular cells and in clusters that resembled proleg epithelium primordia (Fig. 1J, Supp Fig. 5). These results indicate that prolegs and pleuropodia can develop side by side on the same segment, under a partial Hox gene knockout, and are not sharing the same embryonic primordia. *abd-A* is necessary for *Dll* expression in the proleg GRN and is also necessary for repressing *Dll* in the leg/pleuropodia GRN in the A2-A8 abdominal segments.

To further confirm this finding, we focused on the function of *Dll* during proleg development. If prolegs and legs are serial homologues, *Dll* should be necessary for proleg development as it is in legs (*13*). However, no *Dll* crispant embryo showed abnormalities in proleg development, suggesting that *Dll* is not involved in proleg development in *Bicyclus*, as also found in sawfly prolegs (*14*).

Three members of the *Sp* family of transcription factors play important roles in appendage outgrowth and activate *Dll* in the leg network of flies (*15*). To test whether orthologs of s*p* genes were co-expressed with *Dll* in the thoracic legs of *B. anynana*, as well as in prolegs, we isolated two of the three members of the *Sp* family, *sp4* and *sp5*, and looked at their mRNA expression domains (*16*). Both genes were expressed in the thoracic legs and in pleuropodia-like domains in the A1 segment, overlapping Dll expression (Fig. 2). However, both genes were absent from proleg tips, where Dll is normally expressed, and instead overlapped the Abd-A protein expression domain in prolegs (Fig. 2 and Supp Fig. 1). These expression domains suggest that *sp* genes do not drive *Dll* expression in prolegs, as they do in legs, and Dll expression in prolegs belongs to a distinct GRN. Interestingly, in segments A3-A6, both *sp* genes were not expressed in a group of cells lateral to the prolegs, perhaps mapping to the region where ectopic pleuropodia emerge upon *abd-A* knockout. This again suggests the existence of two different embryonic primordia in abdominal segments, one of them expressing *sp4* and *sp5* in circular patches of cells in A1, but not in the same homologous patches in A3-A6, and another expressing *sp4* and *sp5* at the base of the prolegs in the A3-A6 abdominal segments.

**Fig. 2.**
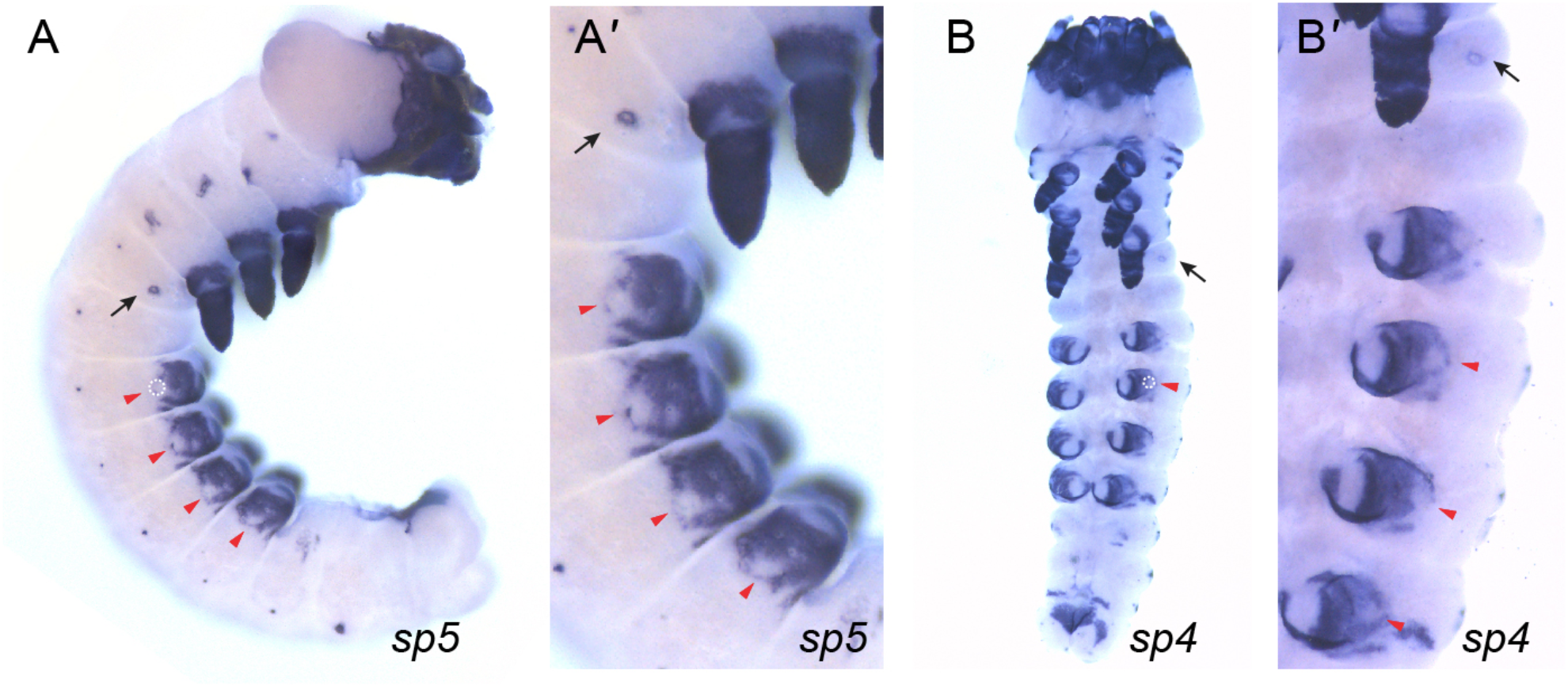
*sp4* and *sp5* mRNAs map to *Dll* expression domains in thoracic legs and pleuropodia but not in the prolegs. Black arrows: pleuropodia, red arrowheads and dotted circles: no *sp4* and *sp5* mRNA expression in cells lateral to the prolegs.

To further examine how proleg and leg GRNs compare to each other and also relative to other appendages in the body, we sampled several 5^th^ instar larval appendages (as in *17*) (Fig. 3A). We extracted total RNA from each tissue, performed pair-wise differential expression (DE) analyses between tissues, and identified 5968 DE genes (logFoldChange ≥ |2| & padj 0.001) across all tissues. Hierarchical clustering (HC) and principal component analyses (PCA) constructed using the DE genes showed prolegs clustered closest to head horns, and formed a sister clade to legs, antennae, and maxillae, with forewings and hindwings forming an outgroup (Fig. 3B and C). To examine whether proleg-specific DE genes also produced the same clustering pattern, we identified proleg-specific DE genes (logFC ≥ |1| & padj 0.05) by comparing prolegs with abdominal regions without prolegs (Supp Fig. 6, Data S1). HC analyses using the identified 2503 proleg DE genes resulted in prolegs clustering on their own to a group containing all other appendages, except wings, the outgroup again in this analysis (Supp Fig. 7A). We also performed HC using 24 leg-specific genes (identified from the *Drosophila* literature) which resulted in prolegs as outliers to all other tissues (Supp Fig. 7B). When we repeated the HC with a subset of 5968 DE genes corresponding to DNA transcription factors and DNA binding proteins (GO:0003700 & GO:0008134), which are the building blocks of GRNs, prolegs and horns clustered together forming a sister clade to all other tissues including the wings (Supp Fig. 7C). Overall, these results suggest that at the 5^th^ instar larval stage, prolegs and head-horn GRNs are most similar.

**Fig. 3.**
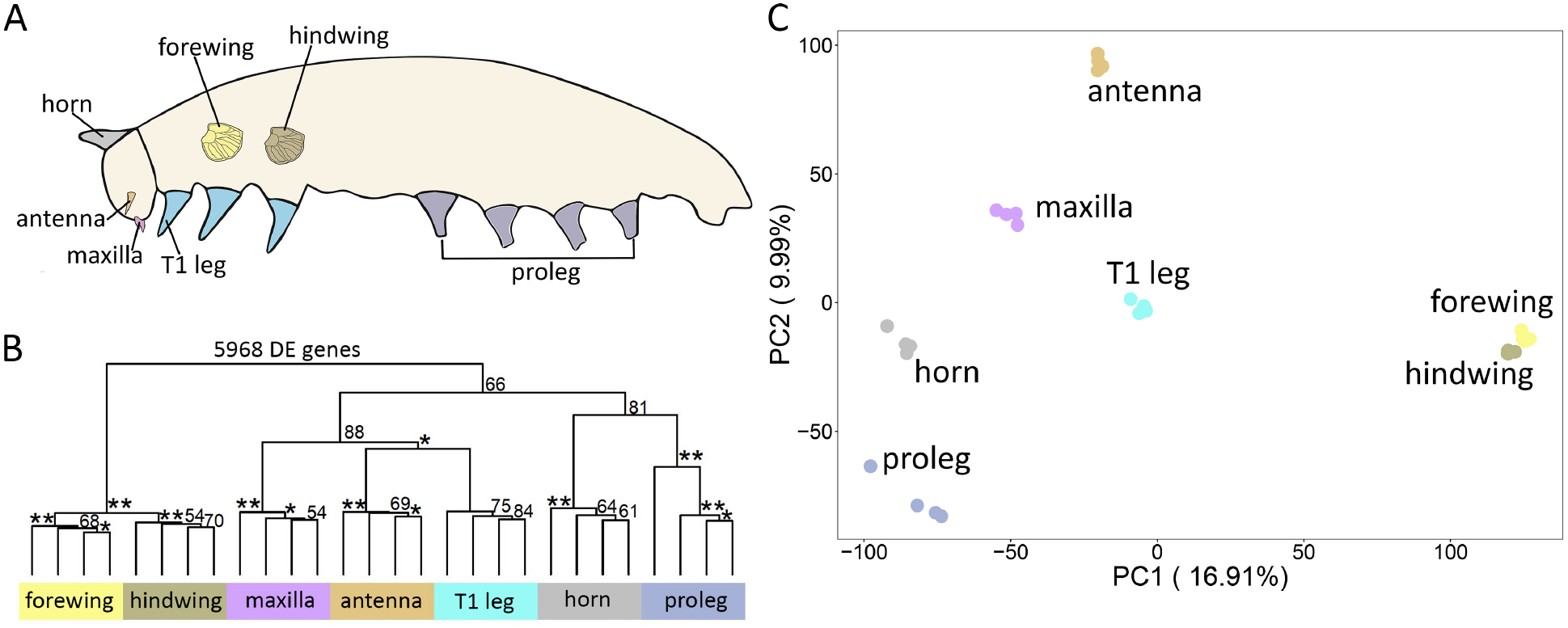
RNA-seq analysis showing proleg transcriptome is closest to head horn. (A) Tissues extracted from the late larval stage for RNA sequencing. (B) Hierarchical clustering using 5968 DE genes from pair-wise comparison between the different tissues. (C) PCA constructed using the variance stabilizing transformation (VST) counts of 5968 DE genes.

In this study, we have shown that *Ubx* and *abd-A* alter the development of two incipient appendages in the abdomen of *B. anynana*, a representative lepidopteran. Ubx is expressed in A1 and modifies the limb GRN there into a pleuropodium (Fig. 4). Abd-A is expressed together with Ubx in A2, where *Ubx* represses the development of the proleg GRN and *abd-A* represses the leg/pleuropodium GRN. In A3-A6, *abd-A* promotes the proleg GRN, and represses the leg/pleuropodium GRN. These results suggest that two traits with different primordia and X-Y coordinates can develop in the same segment and are affected in opposite ways by *Ubx* and *abd-A*.

**Fig. 4.**
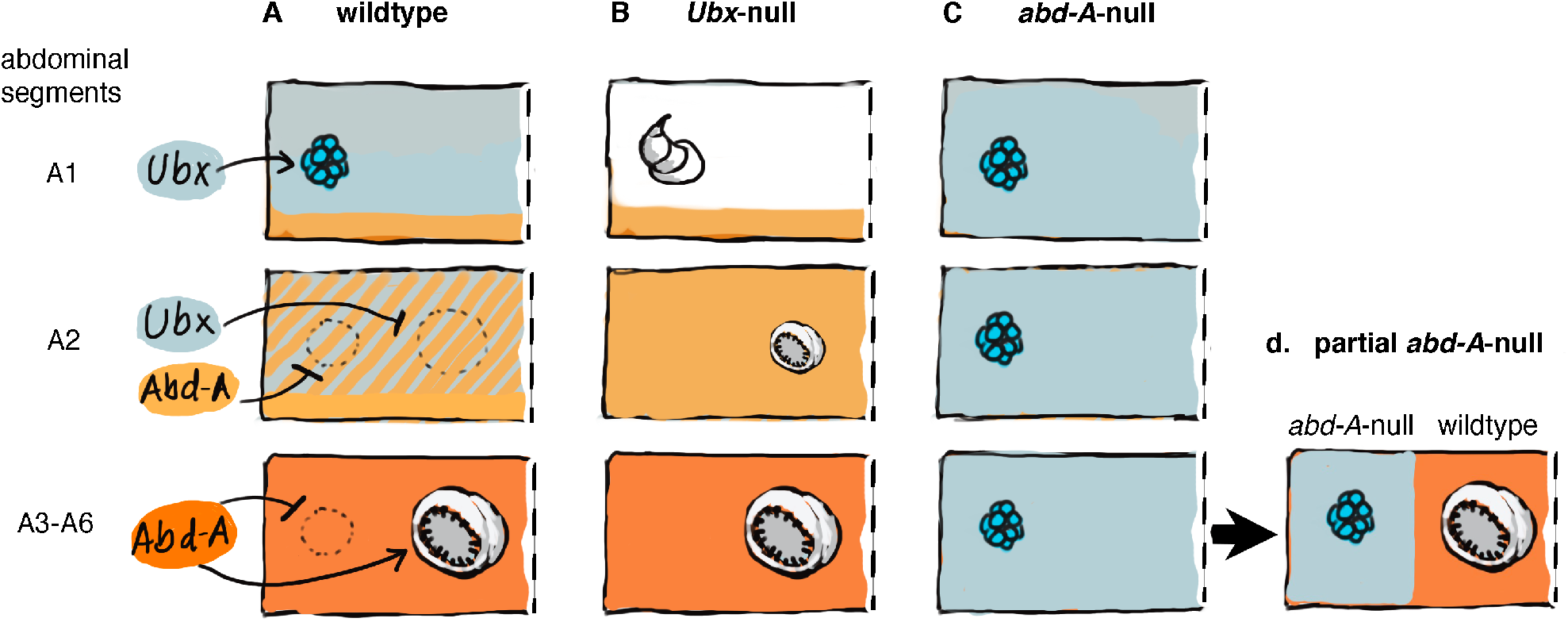
Summary of how null mutations in two Hox genes alter the development of pleuropodia and prolegs in the abdomen of *B. anynana*. Each rectangle represents half a body segment.

Furthermore, we showed that *Dll* and *sp* genes are not co-expressed in prolegs as in thoracic legs, indicating that prolegs have a separate GRN. Previous work had identified a *cis*-regulatory element (CRE) of *Dll* that when expressed in a reporter construct drove EGFP expression in antennae, mouthparts, and thoracic legs, but not in prolegs (*17*). When this CRE was disrupted, all traits were similarly disrupted apart from prolegs. These data suggest, again, that *Dll* expression in the tips of prolegs is driven by a separate CRE, perhaps part of a proleg-specific GRN.

We showed that prolegs and legs/pleuropodia are distinct traits regulated by *Ubx* and *abd-A* in opposite ways, which prevents them from coexisting in the same segment. This result calls for a reexamination of the role of Hox genes in the development of other novelties in the body of insects (*11*). These include gin traps in the abdomen of *Tenebrio* beetles (*18*), gills in the abdomen of mayfly nymphs (*19*), horns in the thorax of dung beetles (*20*), and helmets in the heads of treehoppers (*21*), all previously proposed to be wing serial homologs. Hox genes present in these body regions could be promoting the development of these novel traits while also simultaneously repressing the development of wings. RNAi manipulations cannot address whether both traits can develop in the same segment, as shown here for prolegs and pleuropodia, because the down-regulation of the Hox gene will derepress wings while perhaps failing to promote the novel trait. This manipulation will appear as if one trait is being transformed into the other, when that may not be the case. Partial CRISPR knockouts, however, might be able to uncover whether these traits have separate embryonic coordinates, are under distinct regulation by the same Hox genes, and can develop simultaneously in the same body segment (*11*).

Our data suggest that prolegs and head horns share a similar GRN that may ultimately derive from a partial co-option and modification of the antennae/maxillae/leg GRN. The same co-option mechanism might explain the evolution of sawfly prolegs (*22*) and the evolution of the abdominal tubercules bearing gills of neuropterans (*1*). It remains to be investigated how many times independently this co-option took place, or whether all these lobular abdominal appendages have a single older origin.

## Acknowledgments

We thank Firefly farms and Grenology for corn supplies, L. S. Shashidhara, Shuichiro Tomita, and Grace Boekhoff-Falk for the anti-Ubx, -AbdA, and -Dll antibodies, respectively, and members of the Monteiro lab for support.

## Author contributions

Conceptualization: YM, AM

Methodology: YM, SNM, AP

Investigation: YM, SNM, AP

Visualization: YM, SNM, AP

Funding acquisition: AM

Project administration: AM

Supervision: AM

Writing – original draft: YM, AM

Writing – review & editing: YM, SNM, AP, AM

## Data and materials availability

RNA-seq data and the transcriptome assembly sequences generated are available in NCBI under Bioproject: PRJNA831205.

## List of Supplementary materials

Materials and Methods:

Supplementary Text

Figs. S1 to S7

Tables S1 to S2

References (23 – 35)

Data S1

## Supplementary Materials

### Materials and Methods

#### Butterfly husbandry

*Bicyclus anynana*, originally collected in Malawi, have been reared in the lab since 1988. The caterpillars were fed on young corn plans and adults on mashed banana. *Bicyclus anynana* were reared at 27℃ and 60% humidity in a 12:12 light:dark cycle.

#### Short guide RNA design

Short guide RNA (sgRNA) target sequences were selected based on their GC content (around 60%) and the number of mismatch sequences relative to other sequences in the genome (>3 sites). In addition, we selected target sequences that started with a guanidine for subsequent in vitro transcription by T7 RNA polymerase.

#### sgRNA production

The template for in vitro transcription of sgRNA was made with a PCR method described in *23*. The forward primer contains a T7 RNA polymerase binding site and a sgRNA target site (GAAATTAATACGACTCACTATAGNN_19_GTTT TAGAGCTAGAAATAGC). The reverse primer contains the remainder of sgRNA sequence (AAAAGCACCGACTCGGTGCCACT TTTTCAAGTTGATAACGGACTAGCCTTATTTTAACTTGCTATTTCT AGCTCTAAAAC). PCR was performed with Q5 High-Fidelity DNA Polymerase (NEB) in 100 ml reaction volumes. After checking with gel electrophoresis, the PCR product was purified with the Gene JET PCR Purification Kit (Thermo Fisher). In vitro transcription was performed with T7 RNA polymerase (NEB), using 500 ng of purified PCR product as a template during an overnight reaction. After DNase I treatment to remove the template DNA, the RNA was precipitated with ethanol. The RNA was then suspended in RNase-free water and stored at -80℃.

#### Cas9 mRNA production

pT3TS-nCas9n, a gift from Wenbiao Chen (Addgene plasmid # 46757), was linearized with XbaI (NEB) and purified by phenol/chloroform purification and ethanol precipitation. *In vitro* transcription of mRNA was performed using the mMESSAGEmMACHINE T3 Kit (Ambion). One microgram of linearized plasmid was used as a template, and a poly(A) tail was added to the synthesized mRNA by using the Poly(A) Tailing Kit (Thermo Fisher). The A-tailed RNA was purified by lithium chloride precipitation and then dissolved to RNase-free water and stored at -80℃.

#### Microinjection

Eggs were laid on corn leaves for 30 min. Within 2–3 h after egg laying, sgRNA and Cas9 mRNA were co-injected into embryos. At that stage, the embryo is a syncytium and cell membranes will only appear around 4–5 h after egg laying (*24*). The concentrations of sgRNA and Cas9 are listed in Table 1. Food dye was added to the injection solution for better visualization. The injections were performed while the eggs were submerged in PBS. The injected eggs were incubated at 27℃ in PBS, transferred onto moist cotton the next day, and further incubated at 27℃. The hatched caterpillars were moved to corn leaves and reared at 27℃ with a 12:12 h light:dark cycle and 60% relative humidity.

#### Immunohistochemistry for embryos

Forty eight-hour embryos were dissected in PBS buffer under the microscope. The samples were fixed in 4% formaldehyde/Fix buffer (0.1 M PIPES pH 6.9, 1 mM EGTA pH 6.9, 1.0% Triton x-100, 2 mM MgSO4) for 30min on ice. The samples were washed with 0.02% PBSTx (PBS + Triton x-100) three times, every 10 min, and then dehydrated with a stepwise Methanol/0.02% PBSTx series from 25%, 50%, 75%, to 100%. The samples were kept in -20℃. For immunostaining, the samples were rehydrated with a stepwise Methanol/0.02% PBSTx series from 100%, 75%, 50%, 25%, to 0.02% PBSTx, and then the samples were kept in 5% BSA/PBSTx for 1 h at room temperature as a blocking reaction. The samples were replaced into the 5% BSA/PBSTx with primary antibody and incubated at 4℃ for overnight. We used a rabbit polyclonal anti-Dll (at 1:200, a gift from Grace Boekhoff-Falk, University of Wisconsin, Madison, WI), a mouse monoclonal anti-Antp 4C3 (at 1:200; Developmental Studies Hybridoma Bank), a rabbit anti-*J. coenia* Ubx antibody (at 1:500; a gift from L. Shashidhara), a mouse monoclonal anti-Ubx/abd-A FP6.87 (at 1:5; Developmental Studies Hybridoma Bank), a rat anti-Abd-A (at 1:300; a gift from S. Tomita), a rat anti-Arm (at 1:1000; *25*). The samples were washed with PBSTx, three times, every 10 min. Then, the PBSTx was replaced with 5% BSA/PBSTx as a blocking reaction for 1 h at room temperature, and then replaced with 5% BSA/PBSTx with an appropriate secondary antibody (1:200), and incubated at 4℃ for 2h. The wings were washed three times in every 10min, and the wings were mounted in ProLong Gold mounting media. The images were taken under an Olympus FV3000 microscope.

#### *In situ* hybridization

*sp4* and *sp5* sequences for RNA probe design were amplified from the total cDNA from two day-old embryos using the primers specified in Supplementary Table 1 and cloned into a pGEM-T easy vector (Promega). After sequence analysis, template DNA for *in vitro* transcription of RNA probes were generated by digesting the vector with EcoRⅠ (NEB). Sense and anti-sense DIG-labeled RNA probes were generated with T7 and sp6 RNA polymerase (NEB) and DIG-labeling Mixture (Roche) according to the manufactures’ instructions.

Forty-eight-hour embryos were placed in PBST and a small hole was made in the egg cases using a pin. Embryos were fixed for 20 min in 4% formaldehyde in PBST followed by 3 washes in cold PBST. The embryos were then dissected from the egg cases and incubated in 25 ∼g/mL proteinase K in cold PBST for 7-8 minutes. They were then washed in 2 mg/mL glycine in cold PBST followed by 5 washes in cold PBST before being gradually transferred to a prehybridization buffer (5X saline sodium citrate (pH 4.5), 50% formamide, 0.1% Tween20 and 100 ∼g/mL denatured salmon sperm DNA). Embryos in prehybridization buffer were incubated at 60°C for 1 hour then transferred to a hybridization buffer (prehybridization buffer with 1 mg/ mL glycine and 180 ng/ml DIG labelled riboprobe) for 24 hours. Embryos were then washed 5-6 times in prehybridization buffer at 60°C, transferred back to PBST at room temperature, washed 3 times in PBST and blocked overnight at 4°C (PBST + 1% BSA). Probes were detected by incubation in 1:3000 anti-DIG alkaline phosphatase (Roche) in block buffer for 1 hour followed by 2 washes in alkaline phosphatase buffer (100 mM Tris (pH 9.5), 100 mM NaCl, 5 mM MgCl2, 0.1% Tween) and a final incubation at room temperature in NBT/BCIP solution (Promega) till color developed. The reaction was stopped by washing in PBST. To mount the thick embryos on slides, slides were prepared by layering parafilm strips to give some height for the cover slips and embryos were imaged on a Leica DMS 1000 microscope.

#### Sample collection and library preparation for RNA sequencing

To examine the transcription profiles of prolegs and other tissues, we extracted RNA from T1 legs, antennae, maxillae (mouthparts), abdomen (proleg-control tissue), prolegs, horns, forewings and hindwings from late 5^th^ instar larvae as described in *17*. We performed the experiment with four biological replicates per group with 10 to 20 individuals in each replicat**e**. RNA was extracted using Qiagen RNA Plus Mini Kit. 25 million reads per samples were sequenced with an average insert size of 250-350bp using Novoseq 6000 and 150bp read length. Library preparation and sequencing was performed at Genewiz, China.

#### RNA-seq transcriptome and annotation

RNA-seq analysis were performed as described in Murugesan et al. 2022. The reads were trimmed and filtered for quality using bbmap tools (*26*). Processed reads were mapped to Bicyclus anynana genome (BaGv2) using hisat2. Stringtie (*27, 28*) was used to produce the combined transcriptome with all libraries using the input from hisat2 output. The final transcriptome assembled resulted in 22689 genes with 39534 transcripts. The transcriptome was annotated using ENTAP pipeline (*29*).

#### Differential Expression (DE) Analysis and Hierarchical Clustering

DE analysis was carried out using the genes read count obtained from Stringtie. To verify if prolegs showed a similar expression profile to legs and its serial homologs, pairwise comparisons were performed between all the tissues, except proleg-control tissue. DE (logFC ≥ 2 & padj ≤ 0.001) genes obtained from the pairwise comparison was used to perform hierarchical clustering (HC) using the **run_DE_analysis.pl** script from Trinity pipeline (*30*). To identify the transcription factors and DNA binding proteins, gene ontology (GO) obtained from ENTAP annotation was used. Proleg specific DE genes (logFC ≥ 1 & padj ≤ 0.05) were obtained by DE analysis between proleg and abdomen region using DESeq2 (*31*) (Data S1). To explore the relationship between prolegs and leg serial homologs using “leg-specific” genes, we produced a list of 38 genes which are known to be expressed and involved in the development of legs using the *Drosophila* literature (Data S1). We obtained the corresponding orthologs in our current *B. anynana* transcriptome using reciprocal BLAST. We rerun the pairwise DE analysis and 24 out of the 38 genes were differentially expressed in at least one of the pairwise comparisons, which we used to perform HC.

### Supplementary Text

#### *Antp* is necessary for the development of proximal regions of thoracic legs

*Antp* crispants had short thoracic legs, lacking a tibia and a femur, showing an intermediate morphology between antennae (Supp Fig. 2B) and thoracic legs (Supp Fig. 2C). They had an unbent claw at the tip, resembling thoracic legs, but they lacked their medial segments, resembling antennae (Supp Fig. 2E and 2F). These results suggest that *Antp* is required to generate tibia and femur segments in *B. anynana* thoracic legs. The role of this gene in prolegs, however, is still unclear as prolegs were not visibly affected in our crispants

Our functional examination of the role of *Antp* in the development of the larval body plan of *B. anynana* showed a slightly different function for this gene, as compared to its function in the lepidopteran *B. mori. Antp* is primarily expressed in the thoracic segments in most insects and provides thoracic appendages their identity (*32, 33*). In Lepidoptera, however, *Antp* is also expressed in a ring of cells partially overlapping with the expression of Dll in the prolegs, which develop in the A3-A6 and in the A10 abdominal segments (Supp Fig. 1A; *12*). We showed that *Antp* is required for the development of the medial segments of thoracic legs, as *Antp* crispants had short thoracic legs, lacking a tibia and a femur (Supp Fig. 2F). The role of this gene in prolegs, however, is still unclear as prolegs were not visibly affected in our crispants. In *B. mori*, RNAi against *Antp* caused fusion of thoracic segments and defects in thoracic legs (*34*), but an *Antp* (*Nc*) *B. mori* mutant showed almost complete transformation of prothoracic legs into antennae and a milder but similar type of changes in mesothoracic legs (*35*). In *Drosophila, Antp* mutants showed partial transformation of thoracic legs into antennae, although the distal segments developed normally (*32*). In addition, *Antp*-RNAi phenotypes in milkweed bug, *Oncopeltus fasciatus*, showed similar truncations of the femur, tibia, and some tarsal segments but still developed normal distal end segments (*33*). It is possible that the requirement of *Antp* to develop both medial and distal leg segments is a *B. mori*-specific function, whereas in *B. anynana, Antp* is showing the more conserved function among insects, which is the development of medial leg segments only.

**Fig. S1.**
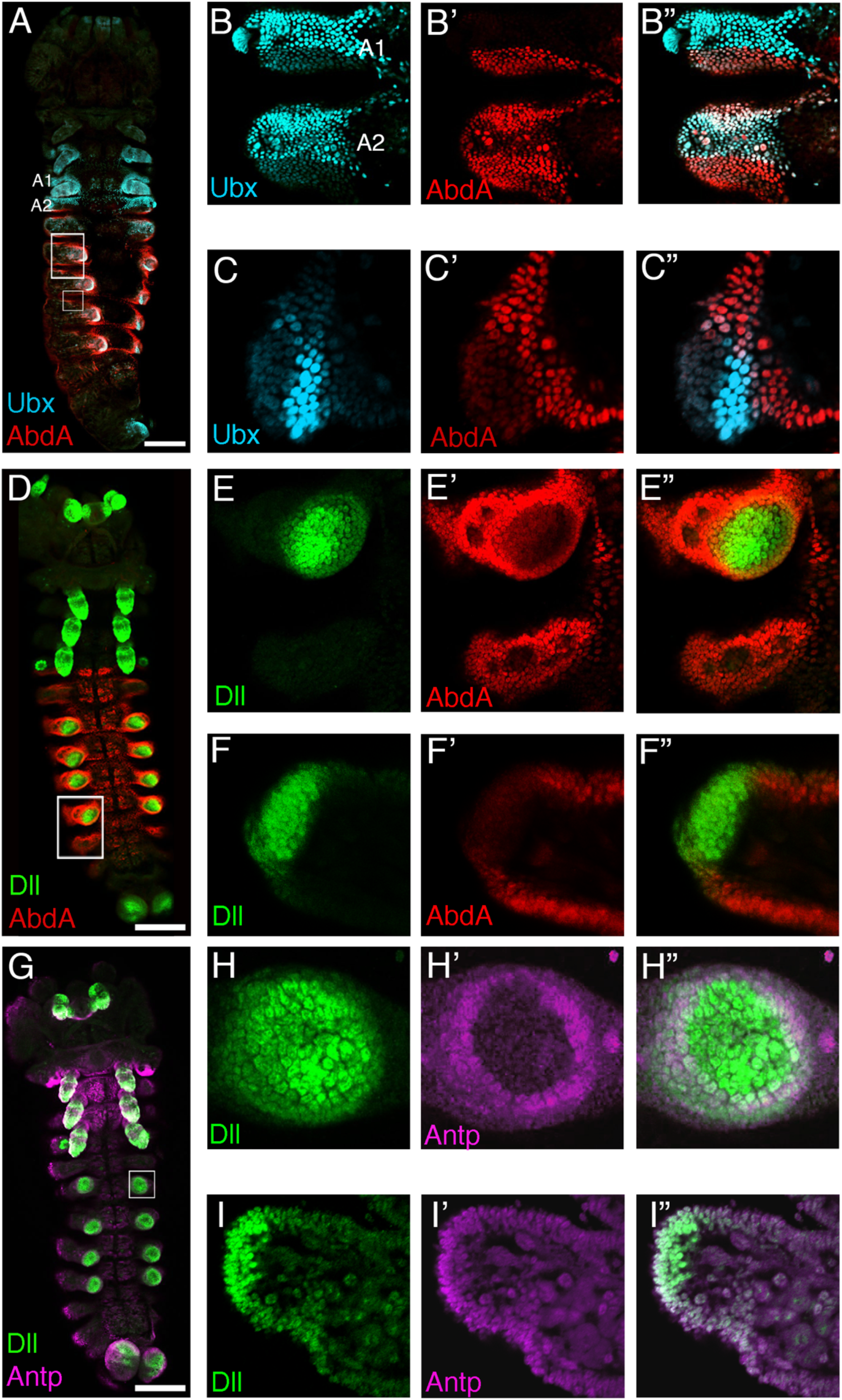
Expression pattern of Ubx, Antp, Abd-A, Dll in wild type embryo. (A) Double staining of Ubx and Abd-A in wild type embryos. Intense expression of Ubx is observed in T2 and T3 legs, pleuropodia, and A1 and A2 segments. Abd-A is expressed in the abdomen. (B) In the A1 segment, Ubx and Abd-A show a clear boundary, while in A2, Abd-A is expressed across the segment, and both genes have overlapping expression domains. (C) In prolegs, Ubx is expressed in the posterior side of the proleg tip. Abd-A is expressed at the base of the proleg. (D) Double staining of Dll and Abd-A in wild type embryo. Dll is expressed in ventral appendages, including antenna, mouthparts, thoracic legs, pleuropodia, and prolegs. (E and F) Dll expression at the tip of prolegs, not overlapping with Abd-A expression. (G) Double staining of Dll and Antp in wild type embryo. Antp is expressed in the thoracic segments and prolegs. (H and I) Antp is expressed at the tip of prolegs, foreshadowing the position where crochets will form. Antp and Dll expression partially overlap. Scale; 100 μm.

**Fig. S2.**
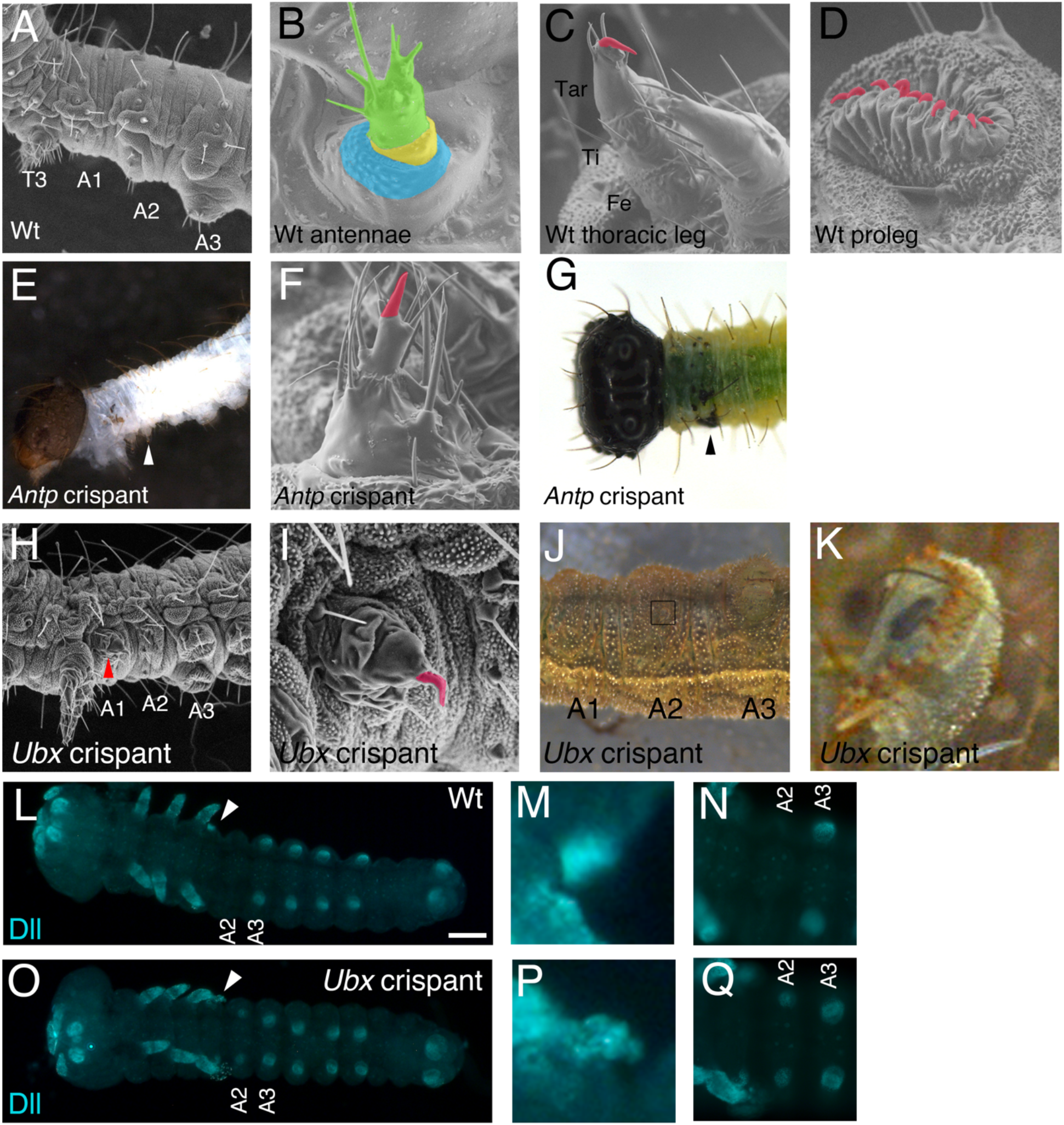
Mosaic phenotype of *Antp* and *Ubx* crispant embryos. (A) SEM picture of a wildtype embryo. (B) SEM picture of a wildtype antenna comprised of three segments, colored in blue, yellow, and green. (C) SEM picture of a wildtype thoracic leg comprised on three segments, femur, tibia, and tarsus. The tibia has setae along its circumference and claw (red) is found at the tip of the tarsus. (D) SEM picture of a wildtype proleg. The proleg has no segments but has crochets at the tip (red). (E) SEM picture of *Antp* mosaic crispant larvae. The T2 and T3 legs of the embryo became smaller compared to the T1 leg. (F) SEM picture of thoracic leg from *Antp* mosaic crispant. The leg has only two segments, and the proximal segment has setae along its circumference. (G) *Antp* mosaic crispant showing dark pigmentation in T2 segment. (H) SEM picture of *Ubx* mosaic crispant larvae. Ectopic protrusion in A1 segment is indicated by red arrowhead. (I) Magnified picture of the ectopic protrusion in A1 segment. The structure likely has no segments but possesses a claw at the tip. (J) Ubx mosaic crispant larva. An ectopic protrusion is found in the A2 segment. (K) Magnified region from the black square in G. The protrusion looks like a small proleg. (L) Expression pattern of Dll in wild type embryo. Dll is expressed in ventral appendages, including pleuropodia (white arrowhead). (M) Magnified picture of Dll expression in pleuropodia. (N) Dll is not expressed in the A2 segment of wild type embryos. (O) Expression pattern of Dll in *Ubx* mosaic crispant embryo. (P) Magnified picture of A1 segment. Dll expression in the protrusion is slightly elongated compared to pleuropodia and resembles a leg (see Fig. S3J). (Q) A2 segment possesses ectopic expression of Dll resembling the pattern observed in prolegs (see Fig. S3K).

**Fig. S3.**
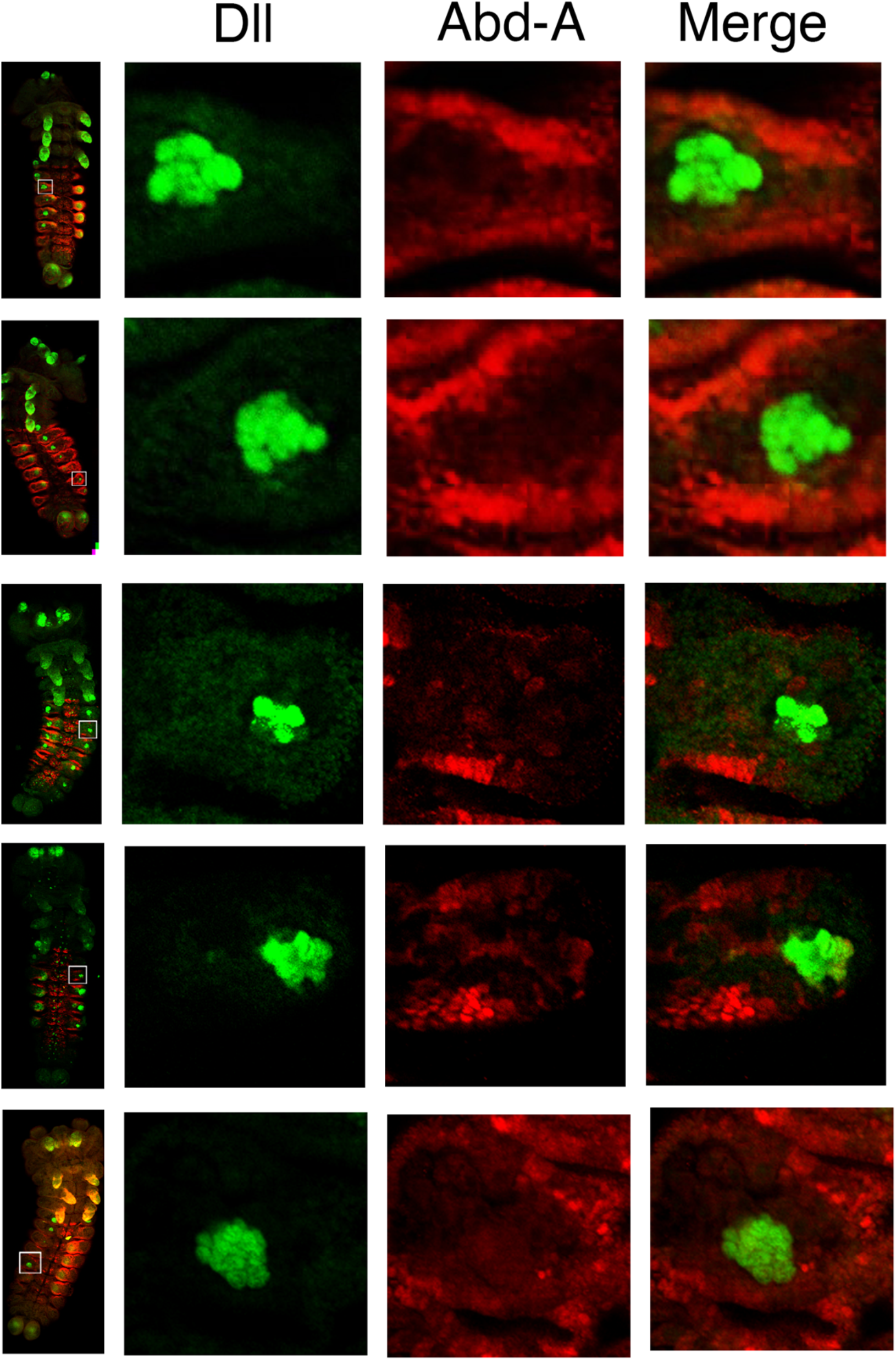
*abd-A* crispant embryos showing loss of Dll expression in prolegs but acquired pleuropodia expression. Thirteen out of 43 *abd-A* crispant larvae loss of Dll expression in prolegs but gain of pleuropodia expression due to the loss of Abd-A activity across the whole segment.

**Fig. S4.**
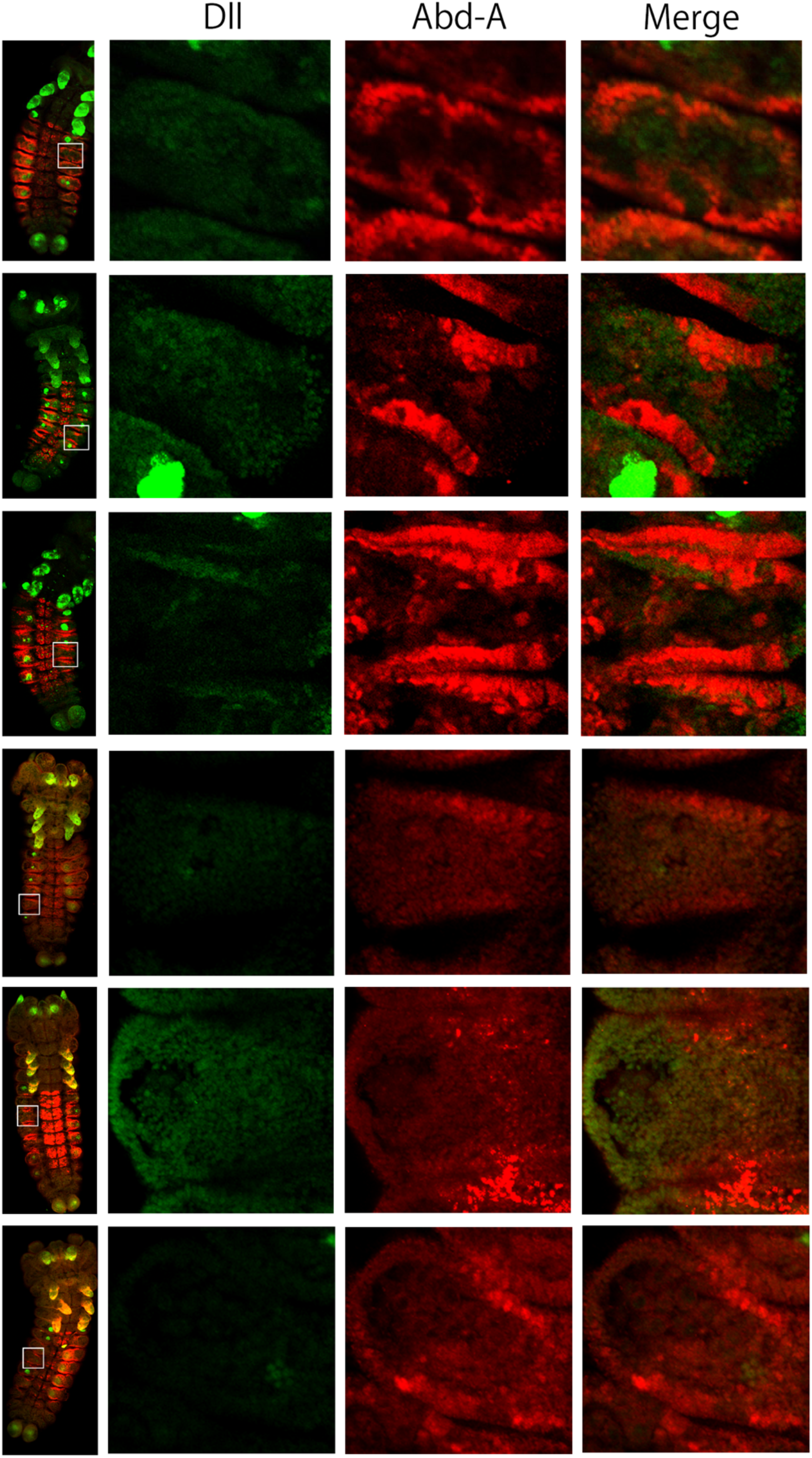
*abd-A* crispant embryos showing loss of Dll expression from prolegs. Twenty out of 43 abd-A crispant larvae lost Dll expression from prolegs due to the loss of Abd-A activity from the mid region of the segment.

**Fig. S5.**
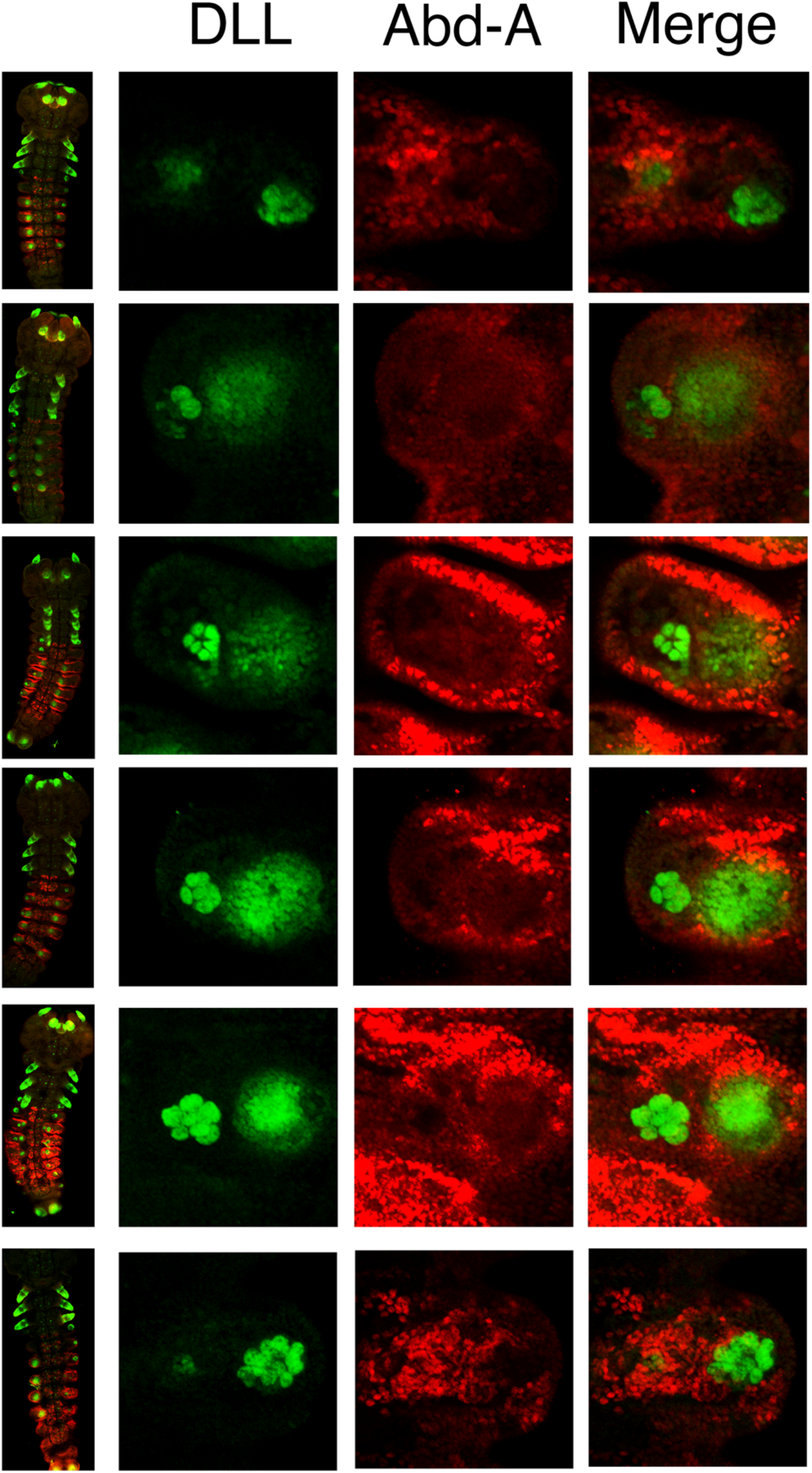
*abd-A* crispant embryos having ectopic pleuropodia. Twenty-eight out of 43 abd-A crispant larvae acquired ectopic pleuropodia due to the loss of Abd-A activity from lateral regions of the segment.

**Fig. S6.**
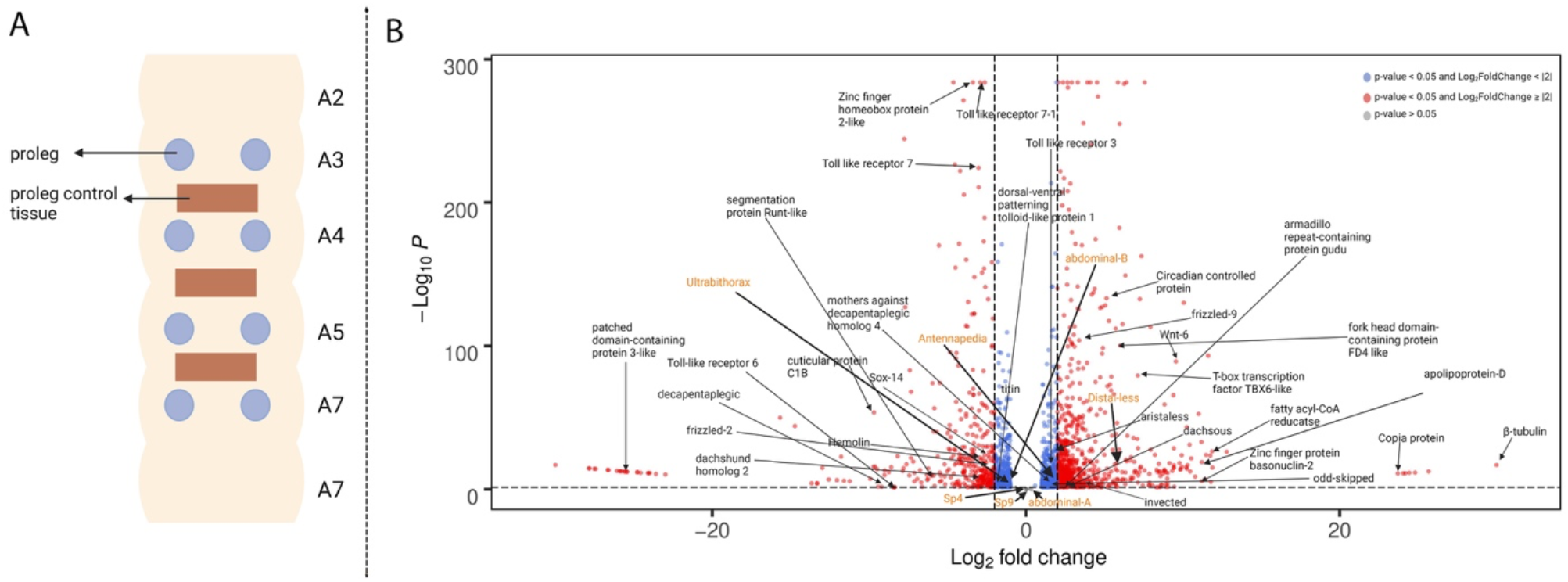
Tissue selection to identify proleg specific DE genes. (A) Abdomen region of *B. anynana* larvae highlighting the prolegs and control tissues in between the abdomen segments used to identify proleg DE genes. (B) DE genes between proleg and control tissues, highlighting the expression level of genes studies in this work in orange color.

**Fig. S7.**
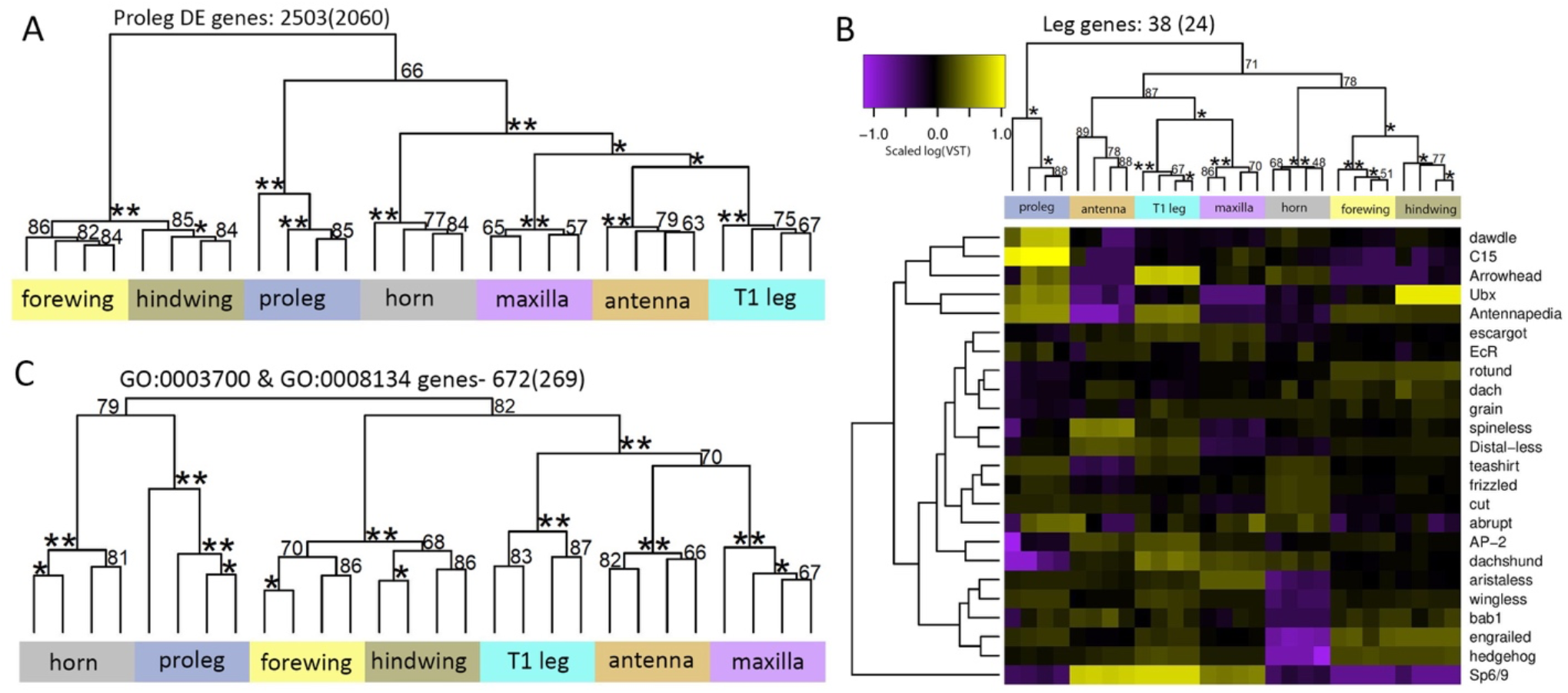
Hierarchical clustering for fifth instar larval tissue using subsets of DE genes shows proleg gene expression is different from that of legs and leg serial homologs. (A) HC using proleg DE genes resulted in proleg clustering as a sister clade to horns, antennae, legs, and maxillae, with forewings and hindwings forming a sister clade to all the other tissues. (B) HC using leg genes shows prolegs as an outgroup to the rest of the larval tissues. (C) HC using DNA transcription factors (TF) and co-factors highlights prolegs and horns share similar expression profile for DNA TF, clustering together and forming a sister clade to the rest of the larval tissues. The numbers mentioned in each HC heading represent the number of genes in each subset. The numbers inside the brackets represent the number of subset genes that were differentially expressed (logFC >|2|, padj 0.001).

**Table S1.**
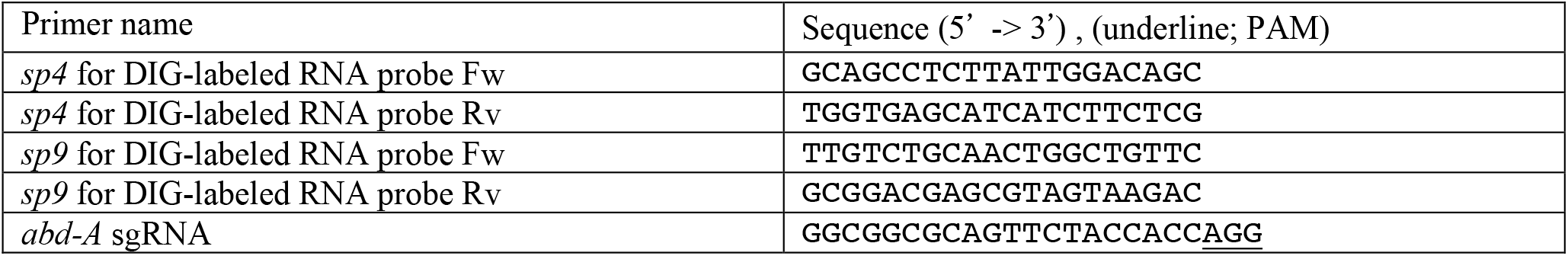
Primer list.

**Table S2.** Summary of CRISPR experiment.

| Total number of abd-A crispant larvae | The crispants showing both loss of proleg and ectopic pleuropodia | The crispants showing only loss of proleg | The crispants showing only ectopic pleuropodia |
| --- | --- | --- | --- |
| 43 | 13 | 20 | 28 |

### Data S1. (separate file)

Proleg DE genes and leg genes orthologs between *D. melanogaster* and *B. anynana*.

